# Host CD59 Potentiates the Type III Secretion System in *Yersinia pseudotuberculosis*

**DOI:** 10.64898/2026.01.07.697569

**Authors:** Kristen J. Davis, Kerri-Lynn Sheahan, Connor S. Murphy, Joshua J. Whiteley, Bree B. Aldridge, Ralph R. Isberg

**Author notes:** Kerri Sheahan, Obsidian Therapeutics; Connor Murphy, Graduate School of Biomedical Sciences and Engineering, University of Maine.

## Abstract

Type III secretion systems (T3SS) play critical roles in disease for many Gram-negative pathogens. Although the structural details of T3SS assembly have been extensively studied, the intimate interactions between T3SS components and the target host plasma membrane are poorly characterized. A previous RNA interference (RNAi) study identified the GPI-linked surface protein CD59 as a critical host component that supports the movement of *Yersinia pseudotuberculosis* (*Yptb*) T3SS effectors into the host cytosol. We show here that both depletion and increased levels of CD59 reduced *Yptb* T3SS pore formation and Yop translocation into host target cells. CD59 is unlikely to act as a receptor for the T3SS translocon, as physical blockade of CD59 or enzymatic removal of all GPI-linked surface proteins had no effect on T3SS function. Consistent with its importance in supporting T3SS function, efficient bacterial effector-mediated focal adhesion disassembly was dependent on CD59. Surprisingly, depletion of CD59 interfered with the rearrangement of the lipid raft marker GM1 that occurs after *Yptb* adhesion to host cells, indicating that CD59 supports lipid microdomain dynamics. Consistent with a role in modulating lipid composition, the loss of CD59 led to the accumulation of phosphatidylcholine and alterations in both fatty acid chain lengths and saturation levels. This work points to a role for CD59 in maintaining plasma membrane dynamics necessary for efficient T3SS translocon function. In addition, a polarization index is described that quantifies the asymmetric association of membrane compartments with organelles.

## Introduction

The pathogenesis of *Yersinia pseudotuberculosis* (*Yptb*) requires the Type III Secretion System (T3SS), which delivers bacterial effectors into host cells, thereby misregulating host signaling pathways and facilitating evasion of innate immune responses (1–3). The T3SS is encoded on a ∼70-kb virulence plasmid present in the three human-pathogenic *Yersinia* species: the enteropathogenic *Y. enterocolitica* and *Y. pseudotuberculosis*, and the causative agent of bubonic plague, *Y. pestis* (4–6). The secretion apparatus is structurally and functionally well conserved among many Gram-negative pathogens, with the *Yersinia* syringe-like nanomachine delivering bacterial effector proteins, called Yops (*<u>Y</u>ersinia* <u>o</u>uter <u>p</u>roteins), into the host cell cytosol (2, 7). There is a core set of six Yops, with YopT absent in most *Yptb* strains (including YPIII, the strain used in this study) (8, 9). Although much work has been done to describe the structure and function of the T3SS, how the host cell supports its function remains largely unknown.

Yops prevent engulfment by professional phagocytes and interfere with reactive oxygen-mediated killing, allowing successful extracellular bacterial replication and survival in host tissues, even with neutrophil-adherent bacteria (3, 10). YopE, for example, is a GTPase activating protein (GAP) that inactivates Rho-family GTPases such as RhoA, Cdc42, and Rac that regulate actin cytoskeletal function required for phagocytosis and production of reactive oxygen species via assembly of the NADPH oxidase complex (11, 12). Interestingly, it has been shown that host cell Rho GTPase activity is initially required for efficient translocation of Yops (13), underscoring that modulation of the actin cytoskeleton plays multiple roles in establishing disease. Several other host factors have been identified that are important for T3SS function, including host cell sulfation and fucosylation in *Vibrio parahaemolyticus*, and DNAJC5 in *Pseudomonas aeruginosa*, which acts as a chaperone for effector translocation (14–16).

We previously identified mammalian proteins important for YopE function using a mammalian siRNA knockdown library and a FRET-based Rho GTPase biosensor (17). The screen identified host proteins, including the chemokine receptor CCR5, which has been shown to affect T3SS function, as well as other proteins involved in cell adhesion, signaling, and membrane trafficking (17). These included CD59, a glycosyl-phosphatidylinositol (GPI)-anchored protein on the surface of most mammalian cell types that functions as a self-recognition marker to protect against complement attack (18). The presence of CD59 on mammalian cells prevents the oligomerization of complement proteins into membrane attack complexes (MACs) upon activation, whereas microorganisms lacking this defense are targeted for destruction (19–21). CD59 is also a direct receptor for some bacterial cholesterol-dependent pore-forming cytolysins and may play a role in forming the T3SS translocation channel (22, 23). In addition, CD59 is an important component of microdomains in the plasma membrane, known as lipid rafts, which are important for critical cell functions such as membrane trafficking, chemokine signaling, and immune cell activation (24). Studies in several similar pathogens, including *Shigella*, EPEC, and *Salmonella,* have revealed that functional lipid rafts on the host cell surface play an important role in supporting microbial adhesion, T3SS translocon pore insertion, and delivery of translocated effector substrates (25–27).

In this study, we show that cells lacking CD59 are associated with aberrant mobilization of host cell plasma membrane components in response to *Yptb*-target cell contact, resulting in defective translocon pore formation and effector translocation.

## Results

### Alterations in surface expression of host cell CD59 result in inefficient pore formation by Yptb

The plasma membrane protein CD59 was identified as a candidate for supporting Yop translocation, based on a pooled shRNA knockdown approach to detect mammalian proteins important for *Yptb* YopE function (17). Verification of the hit revealed that when host cell CD59 was knocked down by shRNA, significant T3SS defects were evident, as assessed by secondary assays for YopB/D-dependent pore formation and YopH translocation (17). These data were consistent with a general defect in T3SS translocator function rather than direct modulation of YopE activity. To pursue these results further, a cloned CD59 knockout HEK 293T line (CD59 KO) was constructed (28). Cell surface labeling of the CD59 KO cell line with anti-CD59-FITC, followed by flow cytometry analysis, resulted in fluorescence indistinguishable from that of an isotype control (Figure 1A). In contrast, WT HEK 293T cells, showed robust labeling with the anti-CD59-FITC conjugate (Figure 1A).

**Figure 1.**
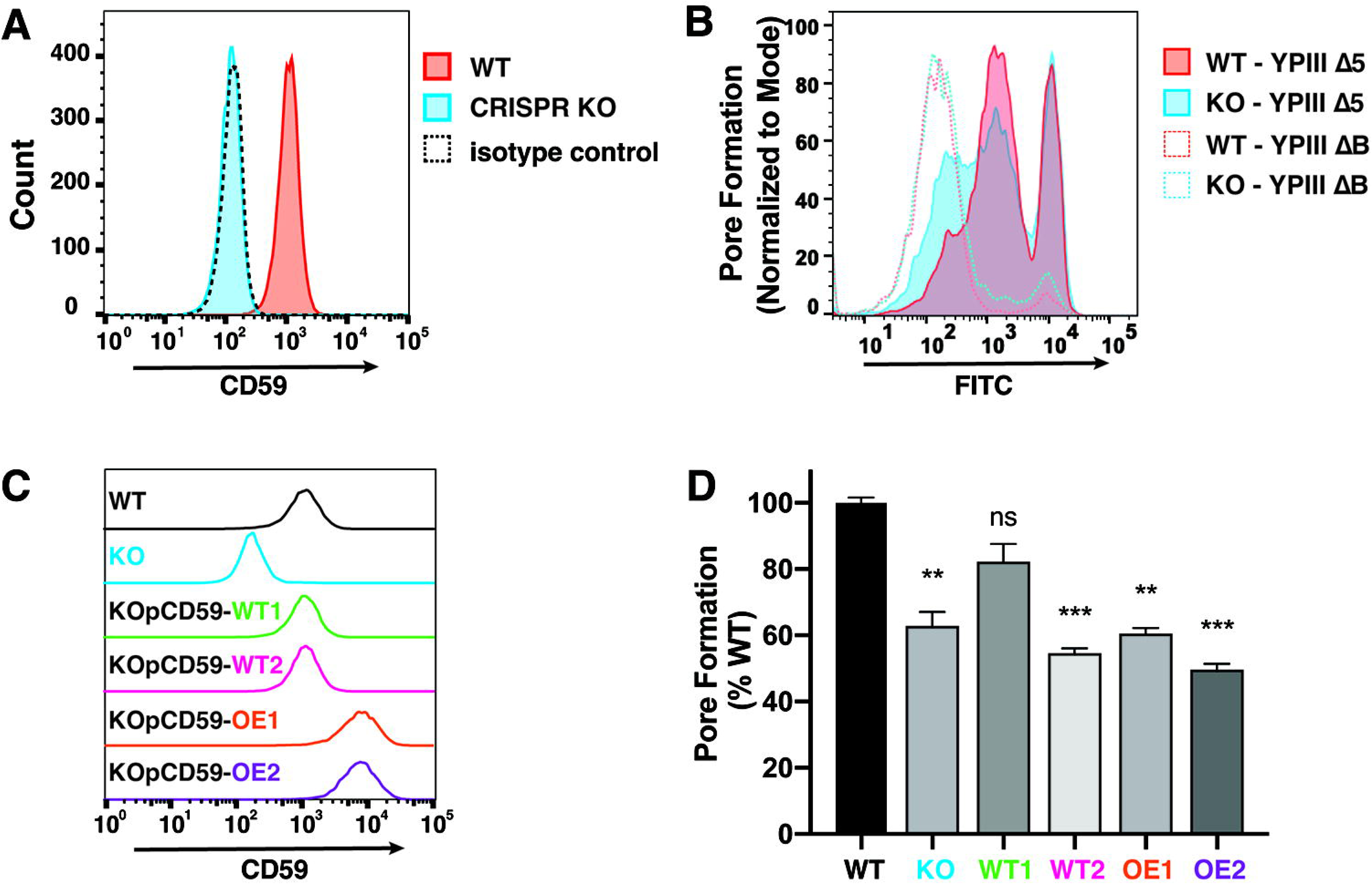
Knockout and increased surface levels of host cell CD59 result in inefficient pore formation when challenged by *Yptb.* (**A)** WT and CD59 KO HEK 293T cells were probed with either FITC-conjugated α-human CD59 or isotype control and analyzed by flow cytometry to quantitate cell surface CD59. Histogram displays total counts for fluorescence of FITC channel, collecting 10,000 events per sample. **(B)** HEK 293T cells were challenged with YPIIIΔ*yopHMOJE* (YPIIIΔ5) or YPIII*yopB* (YPIIIΔB) at MOI = 25 for 2 hours. Cells were labeled with FITC LIVE/DEAD stain and analyzed by flow cytometry to determine extent of pore formation (Materials and Methods). **(C)** CD59 KO 293T cells were transfected with a plasmid expressing human CD59 and stable monoclonal cell lines were obtained with CD59 levels that were similar to endogenously expressed protein (WT1 and WT2) or which were overexpressed (OE1 and OE2). The histogram indicates the level of FITC-conjugated α-human CD59 labeling for the WT, CD59 KO, and pCD59 clones as determined by flow cytometry. **(D)** Indicated cell lines were challenged at MOI = 25 for 2 hrs with YPIIIΔ5 (See Supplemental Figure S2A for YPIIIΔB control). Cells were labeled with FITC LIVE/DEAD stain and analyzed by flow cytometry to determine extent of pore formation. (See Supplemental Figure S2B for pore formation peak data showing WT1, WT2, OE1, and OE2 compared to WT and KO after infection by YPIIIΔ5). Data are geometric mean ± SEM of the fluorescence intensity normalized to WT Results represent three independent experiments performed in technical triplicate. Statistical significance was calculated using a one-way ANOVA with Dunnett’s multiple comparison test (*p < 0.05).

To determine if *Yptb* challenge of the CD59 KO cell line exhibited translocation defects similar to those observed in the shRNA knockdown, a T3SS pore formation assay was performed. Once injected into a target cell, the Yop effectors downmodulate pore formation to prevent immediate cytotoxicity of the host (29). Therefore, a Δ*yopHMOJE* (YPIIIΔ5) strain that lacks all regulatory Yops and exhibits hyper pore-forming activity was used to measure pore formation, measuring fluorescence of an amine-reactive membrane-impermeant probe as a proxy (17).

The *Yersinia* translocon pore complex comprises the two hydrophobic pore proteins YopB and YopD, along with the hydrophilic scaffolding protein LcrV, which localizes to the needle tip (7). A *yopB* mutant strain (YPIIIΔB), lacking the major translocon pore protein YopB, is unable to form pores in the host cell plasma membrane and serves as a negative control for pore formation and translocation (10). WT and CD59 KO cells were challenged with YPIIIΔB or YPIIIΔ5 at MOI = 25 for 2 hours and analyzed for FITC incorporation by flow cytometry. WT and CD59 KO cells infected with the YPIIIΔB mutant were intact and unperturbed, as shown by their similarly low background fluorescence (Figure 1B). In WT cells infected with YPIIIΔ5, two distinct right-shifted populations labeled by the probe were observed. The peak furthest to the right indicates maximum cytotoxicity, while the wider, shouldered peak represents varying degrees of pore formation. The CD59 KO cells also showed two populations, but there was a distinct leftward shift toward lower probe incorporation, indicating that cells lacking CD59 were less susceptible to pore formation by YPIIIΔ5 than WT cells (Figure 1B). Similar results were found using a real-time propidium iodide permeability assay (Supplemental Fig. S1).

To further interrogate the effect of CD59 plasma membrane levels on *Yptb* T3SS function, WT CD59 was introduced into the CD59 KO cell line to evaluate if pore-forming activity could be restored. Human CD59 expressed from the mammalian expression vector pcDNA3.1+ was transfected into CD59 KO 293T cells, and monoclonal populations were isolated after antibiotic selection. Two clones exhibiting CD59 levels similar to WT (KOpCD59-WT1 and KOpCD59-WT2), as well as two clones with increased surface levels of CD59 (KOpCD59-OE1 and KOpCD59-OE2) were identified (Figure 1C). Each of these clones, along with the WT and KO cell lines, was assayed for pore formation after infection by YPIIIΔ5. Interestingly, both CD59 clones with increased surface levels of CD59 showed similar resistance to pore formation by YPIIIΔ5 compared to what was observed with the KO cells. The only clone that restored the degree of pore formation seen in WT cells was one of the clones that exhibited WT levels of surface CD59 (Figure 1D; WT1). Although clones could be isolated that restored pore formation by YPIIIΔ5, there appeared to be confounding effects that prevented effective complementation. In addition to the clones shown here, random screening of several clones that overproduced CD59 relative to WT failed to show the same level of pore formation as the WT1 clone (Figure 1D).

Phenotypic variation was observed among the CD59 KO and KOpCD59 clones after prolonged passaging, introducing another confounding factor. Strategies were sought to overcome this issue. CD59 is one of the most abundant GPI-linked proteins, so knocking it out may select for suppressor mutations or unpredictable changes in regulatory patterns that could be propagated over many passages. For this reason, we continued investigating the role of CD59 in *Yptb* T3SS function using transient siRNA knockdown. Efficient CD59 knockdown was achieved 72 hours post-transfection using mixed pools of siRNAs directed against CD59 (Materials and Methods; Figure 2A).

**Figure 2.**
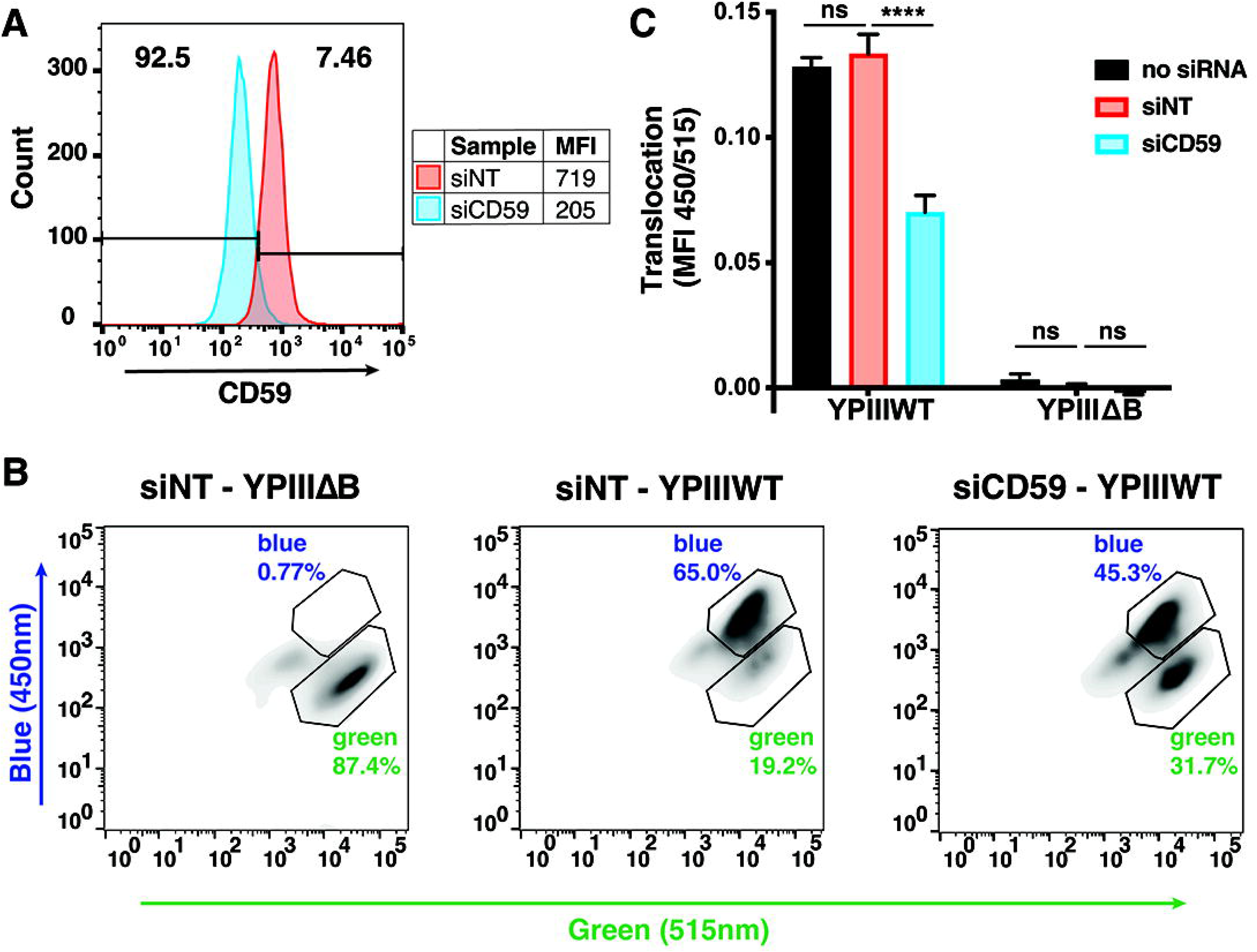
Knockdown of host cell CD59 inhibits translocation of T3SS substrates during *Yptb* infection. WT 293T cells were transfected with either human CD59 siRNA or non-targeting control siRNA to knock down CD59 expression. Assays were performed after 72 total hours of transfection. **(A)** Cells were probed with FITC-conjugated α-human CD59 and analyzed by flow cytometry for cell surface localization of CD59. Bars note the fluorescence distribution of siCD59 sample. **(B, C)** Approximately 2.5 x 10^5^ siRNA transfected cells were seeded overnight. Cells were challenged with YPIII/YopE-TEM or YPIIIΔ*yopB*/YopE-TEM at MOI = 1 for 1 hour. Infected cells were harvested, loaded with CCF4-AM and analyzed by flow cytometry. **(B)** Contour plots displaying the fluorescence intensity of the blue channel (450nm), indicative of YopE translocation, vs. the green channel (515nm) displaying all cells. **(C)** Translocation was calculated as the geometric MFI ratio for the 450/515 channels. Data shown are a representative experiment (from 3 independent experiments performed in technical triplicate). Statistical significance calculated using two-way ANOVA with Dunnett’s multiple comparison test. Not significant: ns; p<0.0001: ****; p<0.05: *.

To evaluate siCD59 knockdown populations using a strategy that does not require loss of membrane integrity or causing cell lethality, T3SS function was assessed by measuring Yop translocation into host cells, detected by assaying for YopE-TEM (or YPIIIΔB YopE-TEM) translocation using a β-lactamase reporter strain followed by FACS analysis (30). After a 1hr infection at MOI = 1, HEK 293T cells were loaded with CCF4-AM, a FRET-based β-lactamase substrate that readily enters the host cells, as detected by a green fluorescence signal at 515 nm after 409 nm excitation. Translocation of YopE-TEM was detected by cleavage of the TEM substrate, which results in a blue fluorescence signal at 450 nm, visualized as the 450 nm emission relative to 515 nm after 409 nm excitation. This was displayed with contour plots of the blue vs. green channels (Figure 2B). Green cells were gated with the YPIIIΔB YopE-TEM negative control (Figure 2B, left panel). After a 1hr infection of siNT cells with YopE-TEM, approximately 65% of the gated blue vs. green cells shifted to the blue gate, indicating robust translocation (Figure 2B, middle panel). In the siCD59 infection (Figure 2B, right panel) this was reduced to 45% of the gated cells, with no detectable translocation in almost 32% of the cells. The total translocation in the population was also reduced by about 50%, as indicated by the ratio of the total fluorescence intensities at 450 nm and 515 nm in siCD59-knockdown cells from triplicate samples (Figure 2C).

### Host cell CD59 indirectly supports *Yptb* T3SS-dependent pore formation

Because pore formation is defective in the CD59 mutant, this protein could act as a receptor for the T3SS channel, so a pair of experiments was performed to test this model. First, a receptor-blockade strategy was employed. Intermedilysin (ILY) is a bacterial-encoded cholesterol-dependent pore-forming toxin (PFT) from *Streptococcus intermedius* that requires CD59 as a receptor (22). The wild-type form of this protein (ILY^WT^) forms pores in the host cell membrane, whereas a mutant version that is fixed in a prepore conformation (ILY^PP^) binds CD59 without forming an active pore, leaving the host cell membrane intact (31). The *Yptb* translocon pore proteins YopB and YopD share physiological properties with PFTs (32), and we hypothesized that if the *Yptb* translocon pore interacts with CD59 similarly to ILY, then ILY^PP^ binding to CD59 will block T3SS-dependent pore formation.

His-ILY^WT^ and His-ILY^PP^ were expressed in *E. coli* BL21 (DE3) and purified (Materials and Methods). HEK 293T cells were pretreated with ILY^PP^ or mock buffer, challenged with ILY^WT^, and assayed for pore formation. Cells that were mock-treated and challenged with ILY^WT^ exhibited high levels of pore formation, confirming that the ILY^WT^ was functioning properly (Figure 3A). Cells pretreated with ILY^PP^ were protected from pore formation, indicating that the mutant protein successfully bound the CD59 receptor and blocked ILY^WT^ (Figure 3A). To determine if ILY^PP^ could block *Yptb* T3SS-dependent pore formation, 293T cells were pretreated with ILY^PP^ for 1 hour and then challenged with YPIIIΔ5 or YPIIIΔB at MOI = 25 for 2 hours (Figure 3B). There was no difference in pore formation among the ILY^PP^ pretreated and mock-treated cells, indicating that if there was a direct interaction between CD59 and the T3SS machinery, it did not occur at the ILY binding site.

**Figure 3.**
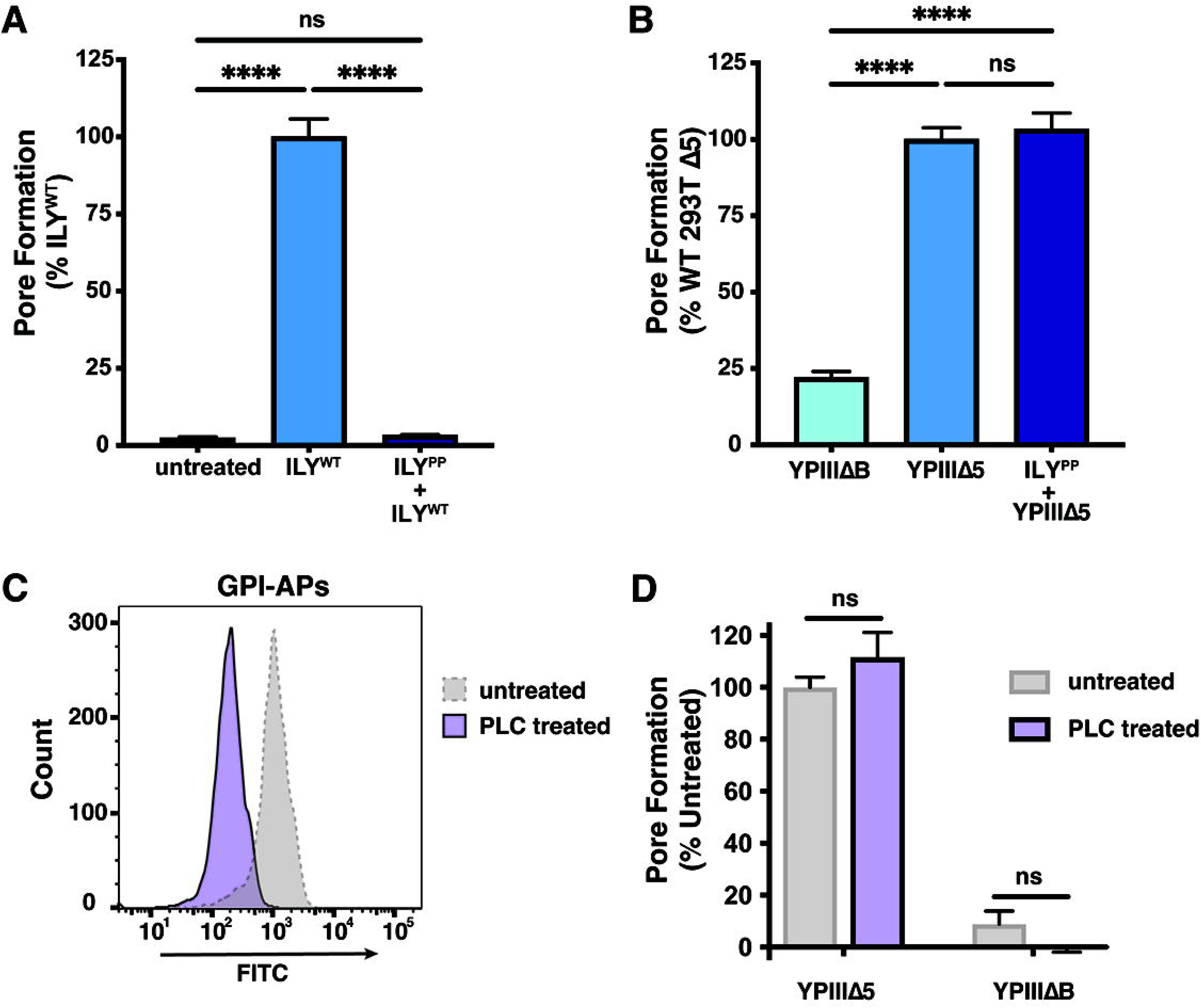
Host cell CD59 indirectly supports YPIII T3SS-dependent pore formation. **(A,B)** ILY^PP^ treatment effectively blocks binding and subsequent pore formation by ILY^WT^. WT 293T cells were seeded overnight in 12-well plates. Cells were pretreated with 100nM of ILY^PP^ or buffer for 1 hour, challenged with either 30nM ILY^WT^ **(A)** or YPIIIΔ5 or YPIIIΔB at MOI=25 **(B)** and incubated for 2 hours. Cells were harvested and assayed for pore formation. Results shown in panels A and B represent the mean from three independent experiments in technical duplicate or triplicate, and all pore formation data were normalized to untreated 293T cells challenged by YPIIIΔ5. **(C)** WT HEK 293T cells were seeded overnight in 12-well plates. Cells were treated with phospholipase C (PLC) and incubated at 37°C, 5% CO_2_ for 30 mins before harvesting and labeling with FLAER for detection of GPI-linked proteins by flow cytometry. **(D)** WT HEK 293T cells were seeded overnight in 12-well plates and pretreated with PLC or buffer prior to challenge with YPIIIΔ5 or YPIIIΔB at MOI = 25 for 2 hrs. Cells were then harvested and assayed for pore formation. Data are geometric means ± SEM of MFI after subtraction of uninfected control cell background and normalized to untreated 293T cells challenged by YPIIIΔ5. Results represent three independent experiments done in technical duplicate or triplicate. Statistical significance was calculated using an unpaired two-tailed Welch’s t test (Not significant: ns; p<0.0001: ****).

A second assay was then performed to determine if CD59 acts as a receptor for the T3SS translocon. To eliminate the possibility that an insufficient quantity of ILY^PP^ was added to the cells to block all binding sites, cells were treated with phospholipase C (PLC), an enzyme that cleaves above the phosphoinositol group of the GPI-anchor, releasing CD59 and other GPI-linked proteins. WT 293T cells were pretreated with PLC, challenged with YPIIIΔ5, then assayed for pore formation. This strategy depleted GPI-linked proteins based on detection of these linkages with Alexa488-conjugated proaerolysin, as PLC treatment resulted in a distinct decrease in fluorescence compared to the mock-treated cell population (Figure 3C). PLC-treated cells were also labeled with FITC-conjugated α-human CD59 to confirm that there was effective removal of CD59 from the cell surface (Supplemental Fig. S3). After the 2 hr infection with YPIIIΔ5, there was no significant difference in pore formation between the PLC-treated and mock-treated cells (Figure 3D). This provides clear evidence that CD59 is not a receptor for T3SS translocation, and additionally rules out any other GPI-linked protein for this potential receptor role.

### CD59 is required for efficient focal adhesion disruption by *Yersinia*

Upon contact with host cells, the bacterial invasin and YadA proteins engage and activate host β1-integrin receptors, leading to recruitment of signaling and cytoskeleton-binding proteins such as paxillin, focal adhesion kinase (FAK), Src-family kinases, and p130Cas. These protein complexes at the integrin cluster sites are called focal adhesions (33). Lipid raft domains are also recruited and involved in integrin clustering and likely promote the assembly and reinforcement of focal adhesions (FAs)(34). *Yersinia* can disrupt these FAs by translocating the YopH protein tyrosine phosphatase, blocking phosphotyrosine signaling and the actin rearrangements that drive phagocytosis (35–37). YopH has many host targets, including FAK, p130Cas, and Src-family kinases at focal adhesion sites (33, 38, 39), as well as the tyrosine kinase Lck and the LAT adapter in T cells (36, 40).

To determine whether CD59 activates focal adhesion formation (40), an anti-phosphotyrosine (pTyr) antibody was used to quantify maintenance of focal adhesions after challenge with the WT *Yptb* strain. To this end, HeLa cells were knocked down with either siCD59 or the siNT control for 72 hours (See Supplemental Fig. S4), challenged with *Yptb gfp^+^* bacteria, fixed, and immunostained with anti-pTyr. Focal adhesions appeared as brightly stained fiber-like bundles, numerous in both siCD59 knockdown and siNT control cells in the absence of infection (Figure 4A). In contrast, upon challenge with WT *Yersinia*, focal adhesions were readily disassembled in CD59-positive siNT control cells, as evidenced by the absence of large bundles and the increased number of puncta (Figure 4B, left panel). This process was inefficient in the siCD59 knockdown cells, as larger focal adhesions were still visible (Figure 4B, right panel) compared to the siNT control cells.

**Figure 4.**
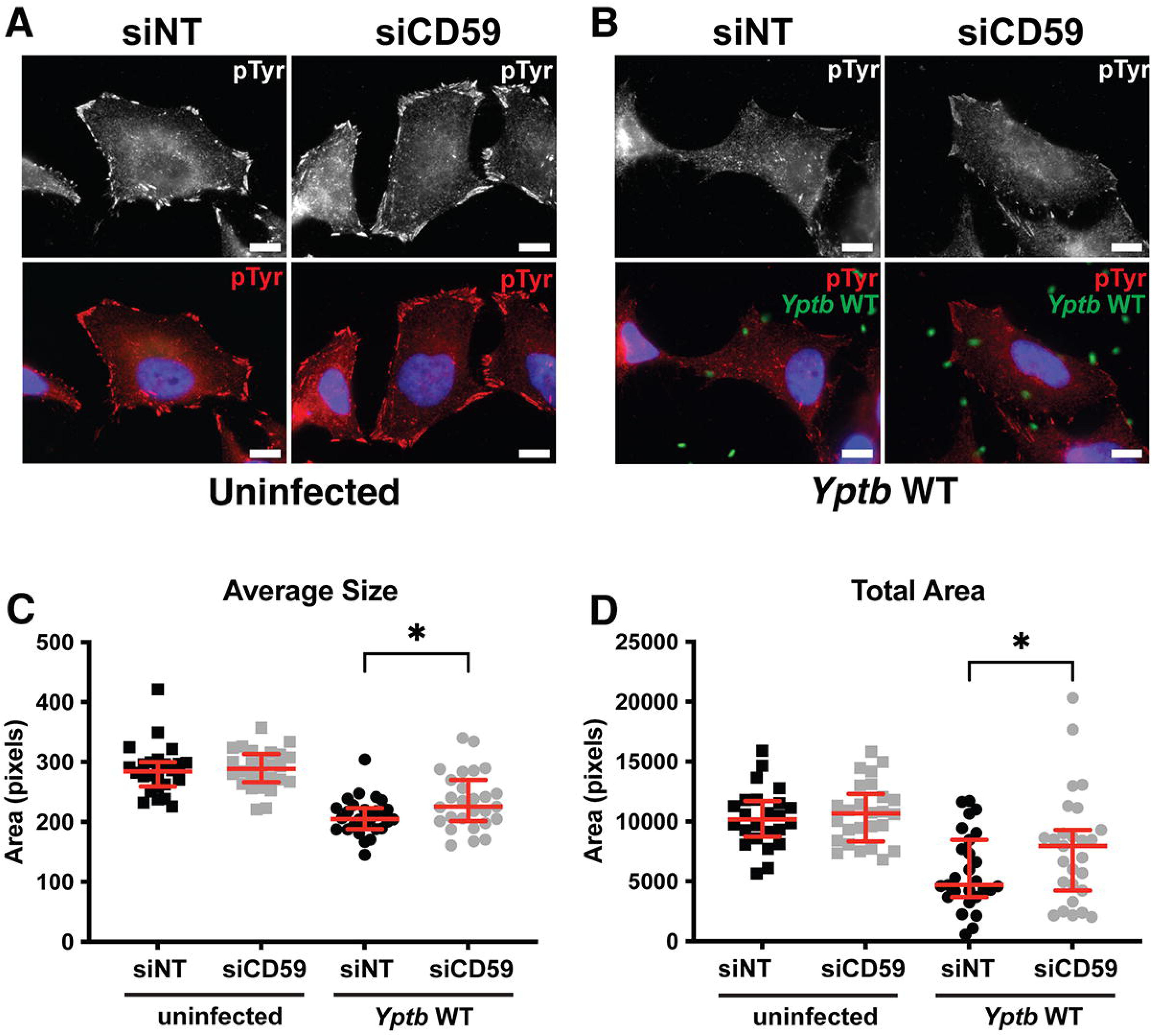
CD59 depletion lowers focal adhesion disruption by *Yptb*. HeLa cells were transfected with either siCD59 or the siNT control for 72 hours. Cells were challenged with *Yptb gfp+* at an MOI = 5 by 5 min centrifugation at 200xg followed by immediate washing and fixing. The cells were then immunostained with anti-a followed by probing with anti-mouse-AF594, and nuclei were stained with Hoechst (1:10000). **(A)** Focal adhesion staining with anti-pTyr of uninfected cells. **(B)** Cells challenged with *Yptb gfp+*. Scale bar = 10μm. Focal adhesions were identified and measured as described (Materials and Methods). **(C, D)** Average size and total area displayed. Data analyzed using Ordinary One Way ANOVA followed by Sidak’s test (*p < 0.05).

Focal adhesions were identified and measured by an image analysis routine that distinguishes edges, based on a previously published workflow (41). The average size of each focal adhesion and the total sum of pixels identified per cell were significantly lower in CD59-positive control cells compared to the CD59 knockdown after bacterial infection (Figure 4C and D, respectively), indicating that CD59 is required for efficient disruption of these host cell signaling assemblies. This is consistent with a defect in YopH accumulation and/or targeting of focal adhesions after bacterial contact with host cells deficient for CD59.

### Depletion of CD59 alters GM1 trafficking after *Yptb* challenge

Our results indicate that the *Yptb* T3SS is sensitive to alterations in cellular CD59 content. CD59 is enriched in microdomains in the cell membrane where cholesterol, glycosphingolipids, GPI-linked proteins, and signaling proteins are densely associated (24). These receptor-rich microdomains are thought to be highly dynamic and may transiently rearrange in response to activating signals, such as pathogen adhesion (42). As CD59 plays an indirect role in supporting *Yptb*-dependent T3SS function (Figure 3), the dynamics of glycosphingolipid GM1 organization were investigated as a function of CD59 expression. Cholera toxin B subunit (CtxB) binds to GM1 in lipid microdomains and can be used to detect altered distribution in response to bacterial binding (43). Therefore, the GM1 probe AlexaFluor594-CtxB was added to the cell culture medium immediately before *Yptb* challenge of cells that had been siRNA-depleted of CD59 for 72 hr prior to infection, using nontargeting siRNA as the control (siCD59 and siNT respectively; See Materials and Methods and Fig. 5G).

**Figure 5.**
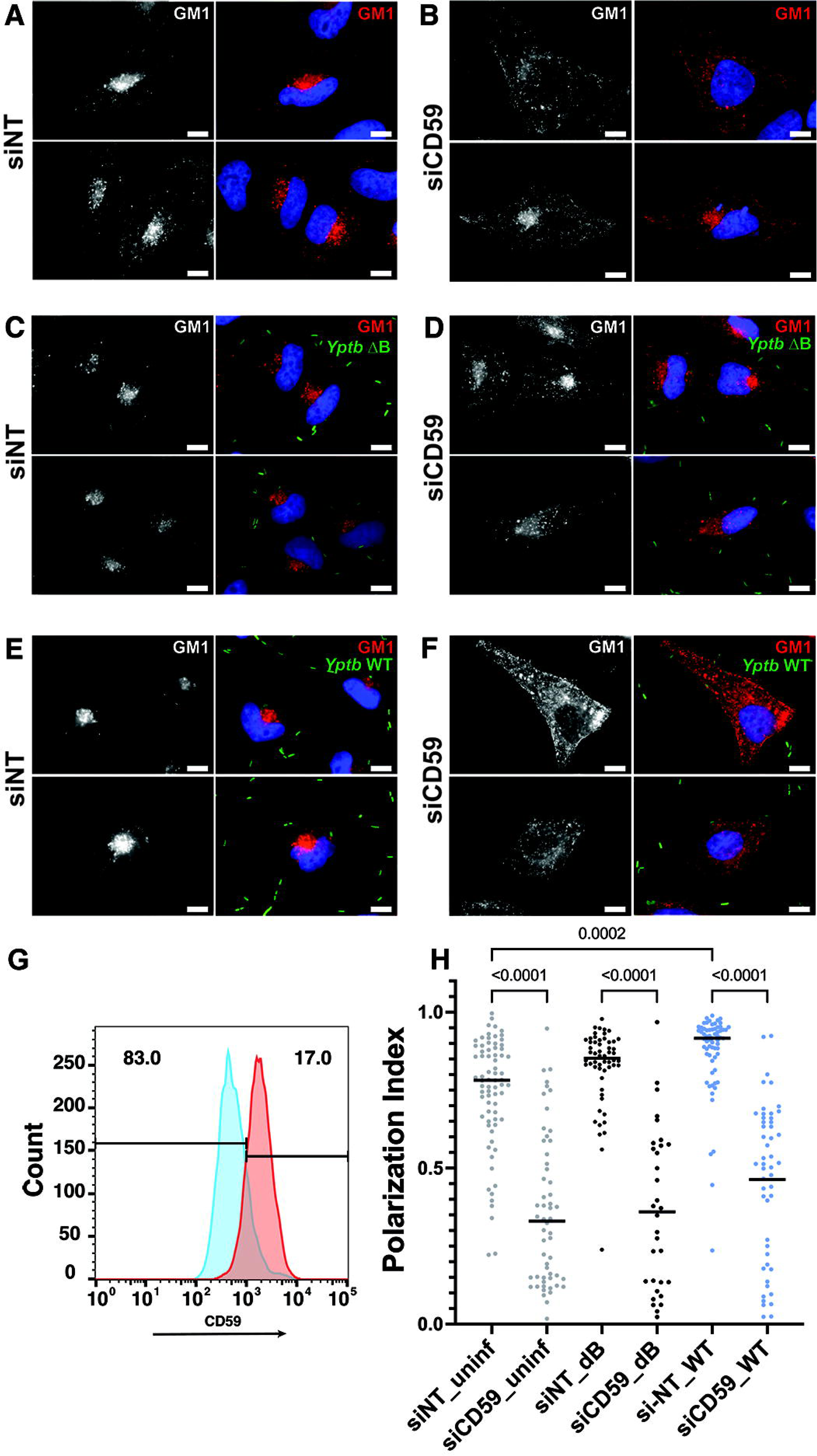
CD59 knockdown affects localization of the lipid raft marker GM1. HeLa cells were transfected for 72 hrs with 25nM of siCD59 or non-targeting siRNA, then seeded on glass coverslips for 24hrs prior to bacterial challenge. **(A, B)** Uninfected. **(C, D)** Cells challenged with YPIIIΔB(pGFP). **(E, F)** Cells challenged with YPIII*-*WT(pGFP). Infections were performed at MOI = 5 in presence of CtxB-AlexaFluor594. Bacterial-host cell contact was initiated by 5 min centrifugation at 200xg followed by immediate washing and incubation with α-Ctx. Cells were fixed with 4% PFA, and nuclei stained with Hoechst (1:10000). Scale bar = 10μm. **(G).** Depletion with siCD59 results in reduced CD59 surface expression compared to control. HeLa cells were depleted with either siCD59 (blue) or siNT (red) for 72 hrs, fixed, probed with anti-CD59 and subjected to flow cytometry to detect surface localization of CD59. Bars note the fluorescence distribution of siCD59 sample. **(H).** Depletion of CD59 blocks polar localization of CtxB proximal to cell nucleus. Polarization Index was determined as described (Materials and Methods; Supplemental Fig. S5) from images of the six conditions described **(A-F)**. Bar: Median of each noted sample. Statistics: Kruskal-Wallis multiple comparisons.

In siNT controls, GM1 was localized in puncta as well as to one side of the perinuclear region (Fig. 5A). After CD59 depletion (Fig. 5G), however, GM1 puncta accumulated at the cell periphery when compared to the siNT control cells, with reduced polar distribution proximal to the nuclei (compare Fig. 5A to 5B, two examples). Challenge with the T3SS-deficient *Yptb* ΔB mutant showed similar results to the uninfected controls (compare Figs 5D to 5B). Strikingly, when siNT cells were challenged with WT *Yptb*, any residual peripheral localization was largely lost and GM1 was almost exclusively localized in a polar fashion proximal to the nucleus (Fig. 5E). In stark contrast, infection of CD59 knockdown cells with WT *Yptb* resulted in little, if any, polar staining about the nucleus, with GM1 puncta dispersed across the whole cell (Fig. 5F), consistent with a defect in GM1 trafficking.

To quantify the CD59-dependent polar staining proximal to the nucleus, an intensity-weighted circular resultant length index was determined for the GM1 localization data (Materials and Methods; Supplemental Fig. 5). To this end, a 1μm wide band about the circumference of each nucleus was determined as a region of interest, with the relative polarization of CtxB localization quantified and expressed as a Polarization Index (R; Fig. S5A-D). Uniform circumferential distribution of CtxB was defined as the Polarization Index R=0, while 100% polar localization was set to R=1.0 (Fig. S5, panel E). Using this index, siCD59 quantitatively blocked polar perinuclear trafficking of GM1 relative to siNT control under all conditions tested (Fig. 5H; P < .0001 for each condition, Kruskal-Wallis multiple comparisons). Also in a quantitative fashion, challenge of cells with YPIII WT increased the polar perinuclear localization of GM1 relative to uninfected controls (P=0.0002), and this trafficking was dependent on the presence of CD59 (Fig. 5H; P<0.0001). Therefore, the absence of CD59 caused the dispersion of GM1 gangliosides and blocked stimulation of polar localization in response to the challenge of WT *Yptb.* This phenotype was not due to alterations in the surface localization of GM1 gangliosides, as flow cytometry of cells probed for GM1 indicated that CD59 knockdown did not affect GM1 surface levels compared to siNT control cells (Figure S6). Therefore, loss of CD59 function resulted in diffuse punctate localization of GM1 and failure of GM1 to traffic to a polar perinuclear locale.

### Loss of CD59 perturbs lipid composition of plasma membrane

To investigate if the loss of CD59 results in changes of lipid composition, HEK 293T WT and CD59-KO cells were extracted in 2:1 chloroform:methanol, and samples were analyzed by LC-MS/MS to identify lipid components (44). Principal component analysis (PCA) of mass spectrometry peak data normalized by percent abundance within each sample showed suitable separation of samples by cell line (Figure 6A). The two most abundant lipid classes in both lines were phosphatidylcholine (PC) and phosphatidylethanolamine (PE). The bulk sum carbon (C) composition of the acyl chains for PC species detected by LC-MS/MS is shown in Figure 6B, indicated by total number of carbons as a percentage sum of all PC lipids for each sample. Acyl chain content species totaling 34 bulk sum carbons was by far the most abundant PC, and the WT had significantly more of these species compared to CD59 KO cells (P<0.05). Interestingly, the KO cells exhibited a greater number of lower (32 carbons) and very large acyl chain content species (>40 bulk sum carbons) (Figure 6B). A similar trend was found in PE species (Figure 6C). WT cells had more of the most abundant 36C content species among PE, while greater numbers of larger (38C) and smaller (<35C) species were detected in CD59KO cells compared to WT cells (Figure 6C).

**Figure 6.**
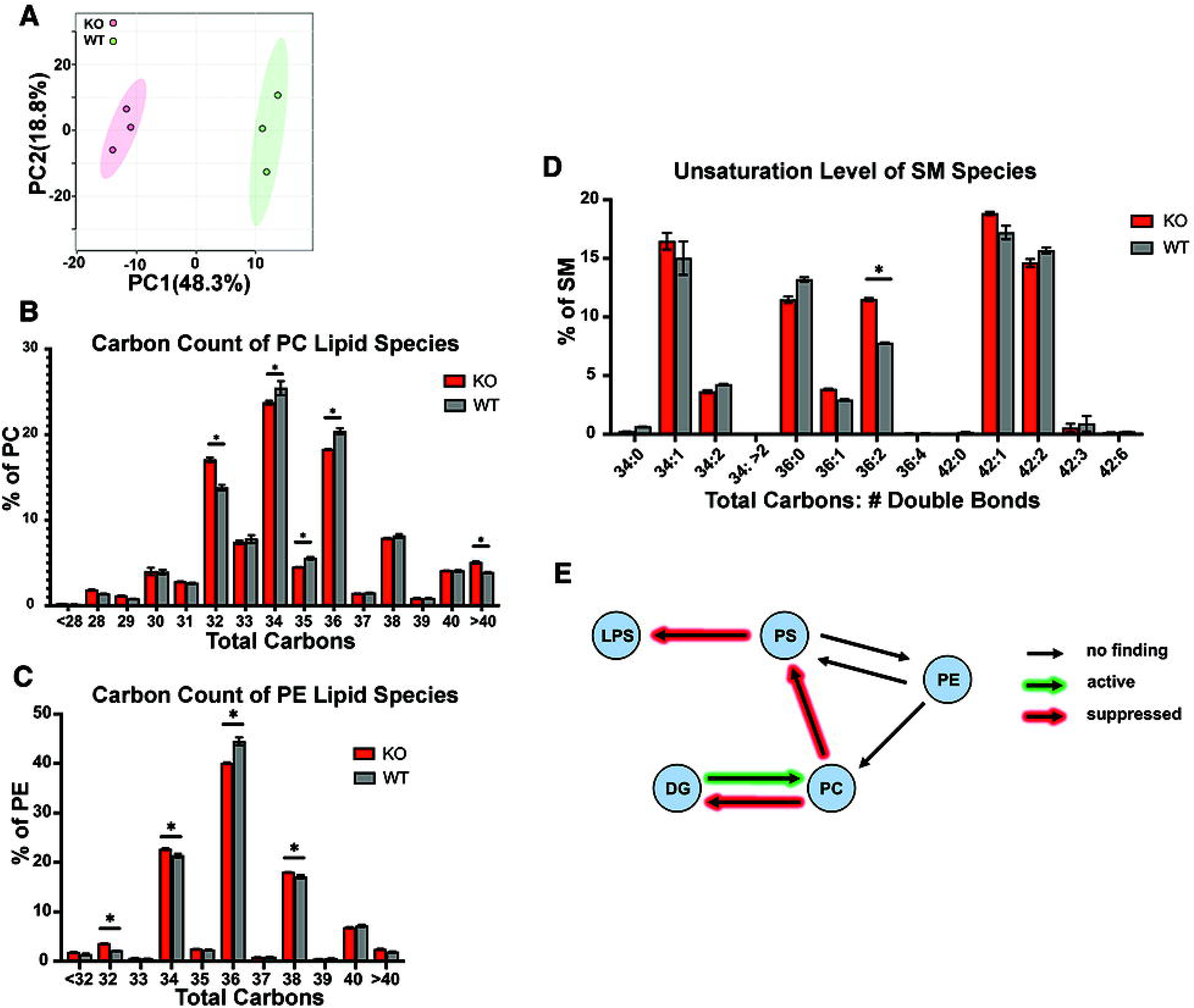
CD59 KO alters cellular lipid composition. Lipids were extracted from HEK 293T WT and CD59 KO cells (Materials and Methods). Representative results of two independent experiments performed in triplicate. **(A)** Knockout of CD59 alters lipid profile. Principal component analysis (PCA) of CD59 KO vs WT cells. Data was normalized by percent abundance within each sample. **(B)** Sum of carbon counts of PC acyl species as a percentage of total PC within each sample. **(C)** Sum of carbon counts of PE acyl species as a percentage of total PE within each sample. **(D)** Saturation levels of SM lipid species as a percentage of total SM within each sample. **(B-D)** Significance determined by unpaired t test using two-stage step-up with FDR of 1% for multiple comparisons, *p<0.000001. **(E)** Alteration of lipid flux as a consequence of loss of CD59. Lipid peak data normalized by percent abundance was analyzed using LIPIDMAPS BioPan software, available through LIPIDMAPS.org (47). Lipid subclass pathways displayed were found to be similarly altered in both experiments. Abbreviations: PC, phosphatidylcholine; PE, phosphatidylethanolamine; SM, sphingomyelin; PS, phosphatidylserine; DG, diacylglycerol; LPS, lysophosphatidylserine.

Sphingomyelin (SM) is enriched in ordered lipid raft domains and has been found to preferentially organize with CD59 within the plasma membrane (45). Increased unsaturation of SM results in greater area and decreased lateral diffusion rates of the lipid (46). There was a significantly higher percentage of SM(36:2) (36 acyl carbons, 2 total double bonds) in CD59 KO compared to WT cells (Figure 6D). The increased unsaturation in the SM 36C species may cuase less-ordered membrane dynamics, potentially affecting lipid raft organization and the ability to coalesce in a perinuclear locale after bacterial contact (Figure 5). CD59 KO also altered flux through several lipid biosynthesis and catabolic pathways, with similar directionality across two independent experiments, as analyzed using the LIPIDMAPS BioPan software tool (Figure 6E) (47). The results of this analysis indicate that in the absence of CD59, there is increased accumulation of phosphatidylcholine (PC) lipids at the expense of phosphatidylserine (PS) and lysophosphatidylserine (LPS) lipids. This should result in membranes becoming less negatively charged in the mutant (PC vs. PS and LPS), while the reduction in lysophosphatidylserine could interfere with trimeric G protein signaling via G protein-coupled receptors (48)(49). Altogether, these results indicate that the absence of CD59 alters the homeostatic maintenance of lipid species composition in the host cell.

## Discussion

In this work, we demonstrated that optimal function of the *Yersinia pseudotuberculosis* T3SS requires CD59. The absence of host CD59 reduced *Yptb* pore formation after cell contact and reduced translocation of effector Yop substrates into target cells. Extracellular blockade of CD59 had no effect on these activities, arguing that CD59 does not act as a receptor for the T3SS. Consistent with an indirect role in supporting the insertion of the translocon channel, the release of all surface-exposed GPI-linked proteins prior to infection did not reduce *Yptb*(Δ5)*-*dependent pore formation. These data, together with the inability of the CD59-binding protein ILY^PP^ to block the T3SS machinery, collectively indicate that CD59 is unlikely to directly interact with pathogen components, but rather supports T3SS translocon assembly at a step after bacterial engagement with the host cell.

Host focal adhesion (FA) complexes are important signaling hubs targeted by the tyrosine phosphatase YopH, which disassembles them and helps prevent bacterial phagocytosis. We found that CD59 facilitates efficient focal adhesion disruption by *Yptb* (Figure 4). In contrast, CD59 depletion increased the average FA size, resulting in a greater total FA area per infected cell. As we previously demonstrated that the absence of CD59 decreases YopH translocation by over 80% relative to control (17), there must be less YopH available to cause focal adhesion disassembly. Alternatively, CD59 has been shown to participate in several signal transduction pathways, including activation of Src-family kinases such as Lyn, Fyn, and Lck, some of which interact with FAK (18, 40, 50). CD59 could be involved in activating or recruiting these components to the site of bacterial binding, so that in its absence, these pathways are muted or less accessible to the Yops. Lipid raft domains, of which CD59 is a hallmark component, are also recruited and involved in integrin clustering and likely promote the assembly and reinforcement of FAs (34). CD59 effects on trafficking may not be limited to host cell components, as its loss may interfere with Yop localization, preventing access to signaling structures normally targeted by the Yops.

A series of studies indicates that the T3SS translocon may interact with lipid raft components. In our previous study, CCR5 was identified as important for T3SS function. CCR5 is in the same family of G protein-coupled receptors (GPCRs) as Fpr1, a recently identified receptor for *Yersinia pestis* that is not required for *Yptb* T3SS function (17, 51). That CCR5 depletion did not completely abolish translocation of *Yptb* T3SS effectors is consistent with *Yptb* targeting functionally redundant receptors, in contrast to the profound importance of Fpr1 for *Y. pestis*. Interestingly, chemokine receptors are known to colocalize with membrane raft markers such as GM1 (52). A similar T3SS-wielding pathogen, *Pseudomonas aeruginosa*, was found to preferentially bind the host receptor glycosphingolipid globotriaosylceramide (Gb3), and depletion of either flotillin or CD59, both found in raft nanodomains with Gb3, reduced uptake of *P. aeruginosa* (53). Functional lipid rafts are important for adhesion and T3SS function in *Shigella*, EPEC, and *Salmonella* as well (25–27). These findings indicate that the organization of various host proteins into structured membrane regions may provide the most efficient target for *Yersinia* to dock, form pores, and deliver Yops via the T3SS.

We examined whether CD59 affected ganglioside-rich domains by fluorescently labeling the raft marker GM1. In uninfected cells, the GM1 marker was predominantly trafficked to the perinuclear region in the presence of CD59. When CD59 was depleted, there were noticeably more GM1 puncta visible around the cell periphery. This phenotype was markedly enhanced upon infection with WT *Yptb.* Accumulation of the GM1 marker was only detected in the perinuclear region in WT host cells, while this perinuclear localization was completely lost in the absence of CD59 after contact with WT *Yptb*. CD59 may play a role in mobilization of GM1 to the perinuclear region either through endocytosis, plasma membrane rearrangements, or recycling of this marker back to the plasma membrane.

When CtxB binds GM1, it is internalized by receptor-mediated endocytosis, and undergoes retrograde trafficking from the plasma membrane through the trans-Golgi network to the ER. This is visualized as perinuclear localization, as we observed in cells with functional CD59 (54). Internalization of GM1 from the surface results in sorting into recycling endosomes followed by release back to the plasma membrane. The first step is visualized as formation of puncta around the cytoplasm, followed by eventual accumulation in pericentriolar recycling endosomes (55, 56). The absence of CD59 may shift how host endocytic pathways are utilized after *Yptb* engagement, favoring a blockade after puncta formation. This could result from changes in the plasma membrane that either prevent effective engagement of the T3SS machinery, alter trafficking of bacterial effectors, or prevent hijacking of host factors by bacterial effectors. Obviously, under CD59 depletion conditions, there must be some low-level or improperly functioning activity of the T3SS that exacerbates the inability of the CtxB-engaged GM1 to coalesce in a perinuclear locale in the absence of CD59.

The specific lipid composition of the host cell plasma membrane can affect its structure, function, and interactions with pathogens. PC and PE are the two most abundant lipid classes. PC is a cylindrical-shaped lipid that allows tight packing of lipids and is present in higher concentrations along with sphingolipids in the outer leaflet of the plasma membrane (57, 58). PE along with phosphatidylserine (PS) and phosphatidylinositol (PI) are mostly found in the inner leaflet (58). Acyl chain length and degree of saturation are lipid characteristics known to affect plasma membrane stability and fluidity (59). Longer acyl chains tend to increase saturation and result in thicker membranes, which are also more resistant to bending (60). Within the PC lipid species, acyl contents of 34C and 36C (34 and 36 total carbons in total acyl chain content) were the most commonly detected acyl chain totals in our analysis. CD59 KO cells had fewer 34C and 36C total chain contents, while exhibiting an increase in longer total acyl chain content (>40) compared to WT cells (Figure 6B). This same trend was observed among the PE lipid species: CD59 KO cells had fewer 36C chain content species, which were most abundant in WT host cells, but had increased very short and very long acyl chain contents compared to WT (Figure 6C).

Acyl chain saturation is also a determinant of plasma membrane organization and fluidity. Fully unsaturated GM1 C16:0 species have been shown previously to co-cluster with CD59, while C16:1 species did not (61). Our analysis showed that SM C36:2 acyl chains were increased in CD59 KO cells compared to WT cells (Figure 6D). The increased levels of unsaturation could have effects on membrane fluidity, raft formation, or lipid-protein interactions. BioPAN analysis revealed altered flux through various catabolic and metabolic pathways, indicating that CD59 is required for proper lipid homeostasis and plasma membrane maintenance (Figure 6E).

In this study, we have shown that CD59 is required for efficient pore formation and translocation of bacterial effectors via the *Yptb* T3SS. The depletion of CD59 revealed changes in the localization of lipid raft marker GM1 indicating a largescale disruption of membrane dynamics. CD59 is an important structural component of raft domains, and it is possible that when cells lack this abundant GPI-linked protein, the formation, stability, and/or mobility of these domains is disrupted. Disruption may result from deranged lipid signaling or from preventing the interaction of T3SS components with host kinases involved in signal transduction. Future work will focus on determining if other host components involved in lipid raft signaling play critical roles in T3SS function, as well as on identifying the critical step in the T3SS translocator insertion process that requires intact CD59.

## Materials and Methods

### Bacterial Strains and Cell Culture Conditions

All *Y. pseudotuberculosis* strains used are derivatives of YPIII (wildtype) and were generously provided by Joan Mecsas (See Table 1). HEK 293T cells (ATCC) and HeLa cells (ATCC) were maintained in Gibco DMEM + 10% fetal bovine serum and incubated at 37°C, 5% CO_2_.

**Table 1.** Bacterial strains used in this study.

| Table 1 Bacterial strains used in this study |  |  |
| --- | --- | --- |
| Background | Strain | Reference |
| YPIII | YPIII$\Delta yopB$ (YPIII$\Delta B$) | (10) |
| YPIII | YPIII$\Delta yopHMOJE$ (YPIII$\Delta 5$) | (10) |
| YPIII | YPIII E-TEM | (66) |
| YPIII | YPIII *gfp*<sup>+</sup> | (67) |
| YPIII | YPIII$\Delta yopB$ *gfp*<sup>+</sup> | (67) |

### T3SS induction

Single colonies of indicated *Yptb* strains were picked from agar plates and grown in 2xYT broth overnight on a rotator at 26°C. Overnight cultures were diluted to A_600_ = 0.200 in induction media (2xYT + 20mM MgCl_2_ + 20mM Na_2_C_2_O_4_), rotated at 26°C for 1.5 hours, then moved to 37°C and rotated for 1.5 hours to induce expression of the T3SS prior to all infection assays.

### Generation of CRISPR CD59 knockout 293T cells

Stable monoclonal CD59 KO HEK 293T cells were generated using the LentiCRISPRv2 vector (a gift from Feng Zhang, Addgene plasmid #52961) (28) targeting the 20bp sequence 5’ ACGACGTCACAACCCGCTTG 3’ of the human CD59 coding region. Oligonucleotides containing this targeted region plus 5’ BsmBI restriction sites (sense: CACCGACGACGTCACAACCCGCTTG and antisense: AAACCAAGCGGGTTGTGACGTCGTC) were annealed and ligated into the LentiCRISPRv2 plasmid at the BsmBI digestion site. The sgRNA-containing vector was linearized within the ampicillin cassette and transfected into WT 293T cells. After 48 hours transfected cells were selected under puromycin and clonal populations were isolated, outgrown, and successful knockout determined using fluorescent antibody labeling and FACS analysis as described below. One knockout clone was used for all subsequent experiments, including the generation of clonal cell lines with wildtype and high levels of CD59 surface expression.

### Generation of CD59 complement and overexpressing 293T cell lines

2 x 10^5^ WT or CD59 KO HEK 293T cells were seeded in 6-well tissue culture-treated plates and the pcDNA3.1 Hygro+ (empty vector) or pcDNA3.1 Hygro+ hCD59 plasmids were transfected using Lipofectamine 2000 (Invitrogen). Hygromycin was added after 48 hours to select for transfected cells, and transfectants were single-cell sorted into 96-well plates containing conditioned cell media. Arising clonal populations were maintained with Hygromycin B to select against loss of the integrated plasmid.

### Translocation Assay

Approximately 2.5 x 10^5^ HEK 293T cells were seeded in tissue culture-treated 24 well dishes overnight. YPIII E-TEM or YPIIIΔ*yopB* E-TEM were grown in induction conditions (See T3SS induction above), diluted in prewarmed DMEM including 10% FBS, and added to 293T cells at an MOI = 1. The infection plates were centrifuged for 5 mins at 1000rpm to maximize bacterial contact with cells and incubated at 37°C, 5% CO_2_, for 1 hr. Cells were then harvested in 180 μl of PBS. A 10X solution of CCF4-AM reagents was prepared in PBS (LiveBLAzer^TM^ FRET - B/G Loading Kit with CCF4-AM, Invitrogen) (62), 20μL was added to each sample tube, briefly vortexed, and incubated for 20 minutes protected from light at room temperature. Samples were then analyzed on a BD FACS Calibur and data was processed with FlowJo (Tree Star) software, with 10,000 events collected for each sample. Translocation was quantified by subtracting background signal present in uninfected cells and calculating the ratio of the geometric mean fluorescence intensity of the 450/515 channels.

### Pore Formation Assay

Approximately 2.5 x 10^5^ HEK 293T cells were seeded in tissue culture-treated 24-well dishes overnight. Overnight bacterial cultures of YPIII Δ*yopHMOJE* (YPIIIΔ5) or YPIIIΔ*yopB* (YPIIIΔB) were grown in T3SS induction conditions prior to infection (See T3SS Induction). Induced YPIIIΔ*yopHMOJE* or YPIIIΔ*yopB* were diluted in prewarmed DMEM + 10% FBS and added to HEK 293T cells at an MOI of 25. After challenge, plates were centrifuged for 5 mins at 200xg and incubated at 37°C, 5% CO_2_ for 2 hours. The contents of each well were removed by pipetting the cells up and down, washing with Hanks’ balanced salt solution (HBSS), Gibco, resuspending in Live/Dead dye diluted 1:1000 in HBSS (LIVE/DEAD™ Fixable Green Dead Cell Stain Kit, Molecular Probes, Life Technologies), and incubated for 30 minutes protected from light. Cells were washed twice in HBSS containing 1% Bovine Serum Albumin (BSA), resuspended in 0.300mL HBSS, and analyzed by flow cytometry on a BD LSRII flow cytometer. At least 10,000 events were collected per sample. Pore formation was quantified by determining the geometric mean of the FITC channel. Data was processed with FlowJo (Tree Star) software.

### CD59 and GPI-linked surface protein detection

Flow cytometry was utilized to determine the levels of CD59 and GPI-linked proteins on the host cell surface. Live HEK 293Ts cells were washed with PBS, blocked on ice in PBS containing 0.5% BSA for 10 minutes, and pelleted for 5 mins at 300rcf. For CD59 detection, cells were resuspended in mouse anti-human CD59-FITC (BD Pharminogen, clone p282 (H19)), diluted 1:500 in blocking buffer. To detect GPI-linked proteins, cells were resuspended in fluorescently labeled proaerolysin (FLAER, Alexa 488 variant, CedarLane Labs) diluted 1:20 in blocking buffer. Cells were incubated on ice protected from light for 30 minutes, washed once in blocking buffer, resuspended, and transferred to a round bottom polystyrene 12 x 75mm tube for flow cytometry analysis. When fixation was required, cells were washed in PBS, fixed for 20 minutes in PBS containing 4% paraformaldehyde (PFA) at RT, protected from light, washed 3x in PBS, blocked for 1hr in 4% BSA, resuspended in either anti-CD59 or FLAER (1:500 or 1:20, respectively) for 1hr rotating at 37°C, washed once, resuspended in blocking buffer, then transferred to round bottom 5mL polystyrene tubes for flow cytometry analysis.

### CD59 siRNA knockdown

HEK 293T or HeLa cells were seeded in 6-well tissue culture-treated plates overnight. 25pmol per well of siGENOME SMARTPOOL human CD59 siRNA (Horizon Discovery) or 25pmol of non-targeting control siRNA were transfected using Lipofectamine RNAiMax (Invitrogen). After 48 hours, cells were lifted and seeded in 24-well tissue culture plates. Cells were then challenged with induced YPIII strains after a total of 72 hours of siRNA knockdown.

### Phospholipase C Treatment

5.0 x 10^5^ HEK 293T cells per well were seeded into 12-well tissue culture-treated plates overnight. Overnight cultures of YPIII Δ*yopHMOJE* or YPIIIΔ*yopB* were then diluted and induced (see T3SS induction). To cleave surface GPI-linked proteins from HEK 293Ts cells, 0.1U of phospholipase C protein (Phosphatidylinositol-Specific from *Bacillus cereus*, Invitrogen), or an equivalent volume of buffer was added to each well 30 minutes prior to infection and incubated at 37°C, 5% CO_2_, then cells were infected at MOI = 25. Plates were centrifuged for 5mins at 200xg and incubated at 37°C, 5% CO_2_ for 2 hours. Following infection, cells were either assayed for pore formation or probed with FLAER to determine levels of GPI-linked proteins present on the host cell surface.

### Expression and purification of intermedilysin proteins

Plasmid constructs for protein expression of His-tagged WT intermedilysin (His-ILY^WT^) and a prepore conformation mutant (His-ILY^PP^) were generously provided by Rodney K. Tweten (63). *E. coli* BL21 (DE3) cultures harboring either the His-ILY^WT^ or His-ILY^PP^ plasmids were grown overnight in LB broth supplemented with 100ug/mL carbenicillin. Overnight cultures were diluted 1:100 in LB and incubated with agitation at 37°C until they reached an A_600_ = 0.8, supplemented with 100ug/mL carbenicillin, and induced by adding 1mM IPTG (final concentration). Cultures were incubated at room temperature overnight and pelleted at 5000rpm for 20 mins at 4°C. Pellets were resuspended in 20mL PBS, 1x EDTA-free cOmplete Protease Inhibitor Cocktail (Sigma), cells were lysed using a microfluidizer (Microfluidics), and the cell lysate centrifuged at 10,000rpm for 30min at 4°C. Cleared lysates were applied to Ni-NTA resin, rotating end over end at 4°C for 1hr to allow adsorption, and mixture was transferred to a plastic column. After allowing flow-through of the adsorption mix, the resin was washed with PBS containing steps of 0mM, 20mM, 50mM, 80mM, and 120mM imidazole, followed by elution of the protein with PBS containing 500mM imidazole, pH =7.2. Protein purity was determined by SDS-PAGE gel and Coomassie staining, and protein concentrations were determined using Bradford reagent (Bio-Rad). Protein was dialyzed overnight into 10mM MES (pH=6.5), 300mM NaCl, 1mM EDTA using dialysis cassettes (Slide-a-Lyzer Cassettes, Thermo), then either stored short-term at 4°C, or long-term by adding 10% glycerol and 5mM DTT before flash freezing for storage at -80°C.

### ILY Treatment

Approximately 5 x 10^5^ WT 293T cells were seeded overnight in 12 well plates. To perform inhibitor experiments, cells were pretreated with 100nM ILY^PP^ or an equivalent volume of buffer per well and incubated at 37°C, 5% CO_2_ for 1 hr. After pretreatment, cells were challenged with 30nM of ILY^WT^ or buffer, incubated for an additional 2 hours, and assayed for pore formation (See Pore Formation Assay).

### Focal Adhesion Immunofluorescence Staining

HeLa cells were seeded on glass coverslips overnight in 24-well plates. Cells were wash and then challenged at MOI = 5, centrifuged for 5 mins at 200xg, then immediately washed and fixed in 4% PFA. The fixed cells were then washed with 0.05 M Tris buffered saline pH 8.0 (TBS) permeabilized in TBS containing 0.1% Triton X-100 for 2-5 mins and blocked with TBS containing 1% BSA. Cells were incubated with anti-phosphotyrosine, clone 4G10 (EMD Millipore) (1:200 in blocking buffer), washed 3 times, and incubated with donkey anti-mouse IgG-Alexa-Fluor 594 (Jackson ImmunoResearch Laboratories). Coverslips were stained with Hoechst, washed, and mounted with ProLong Glass Antifade (Invitrogen). Images were taken with Zeiss AxioObserver.Z1 fluorescent microscope. Focal adhesions were identified and measured in ImageJ using a custom macro that automates various functions to minimize background, increase contrast, and enhance edge detection to identify focal adhesion particles, based on a previously published workflow (41). See Supplementary materials for macro details.

### GM1 Immunofluorescence Staining

HeLa cells were seeded overnight in DMEM containing 10% fetal bovine serum on glass coverslips in 24-well plates. Prior to infection, the culture medium was aspirated and fresh culture medium containing cholera toxin B (CtxB) (Vybrant Lipid Raft Labeling Kit, Molecular Probes) was added. Cells were then infected at MOI = 5 with *Yptb* diluted in DMEM + 10% FBS so that the final concentration of CtxB was 2μg/mL in each well. Plates were placed in centrifuge, subjected to 5min spin at 200xg, then coverslips were washed three times with cold medium, incubated with anti-CtxB at 4°C for 15mins, washed three times with cold PBS, fixed in 4% PBS containing 4% PFA, washed three times in PBS, stained with Hoechst (1:10000) and mounted with ProLong Glass Antifade (Invitrogen).

### Quantification of Polarization Index for CtxB localization

To measure asymmetric localization of AlexaFluor594-CtxB, multichannel 63X CZI images were captured from an Axio Observer Z1 inverted microscope, using Hoescht staining to perform nuclear segmentation, and AlexaFluor594 to quantify perinuclear receptor aggregation of CtxB. Nuclei were first segmented using a Python script incorporating Cellpose-SAM (64) with segmentation masks being cached per field, followed by inspection using Hoescht/mask overlay images (Supplemental Fig. S5D). Nuclei within 20 pixels of the image border were excluded from statistical comparisons because perinuclear-localizing fluorescence could be partially outside the field of view. Once nuclear segmentation was completed, a Polarization Index (R) was calculated by taking each segmented nucleus and defining the associated perinuclear AlexaFluor594 by thresholding the AlexaFluor594 channel with Otsu’s method (65), excluding pixels inside the nuclei, and assigning extranuclear pixels above the Otsu threshold to the nearest nucleus by Euclidean distance transform. The assigned CtxB regions were restricted to a 1.0 μm perinuclear shell. The Polarization Index was then quantified as the intensity-weighted circular resultant length (R), of the assigned mCherry pixels around the nuclear centroid. To this end, a routine was employed to perform these tasks as follows.

For each nucleus, the code segmented the nuclear Hoescht signal, found the above-background perinuclear AlexaFluor594 pixels that defined CtxB localization, then assigned those pixels to the nearest nucleus, computing a one-sidedness score (R).

To calculate **R**, for the AlexaFluor594 channel pixel ***i***, let ***w***□ be its intensity and θ□ be its angle around the nucleus center:

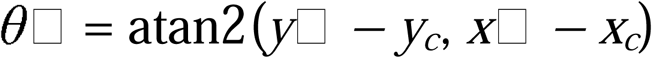

The asymmetry score was then determined as the intensity-weighted circular 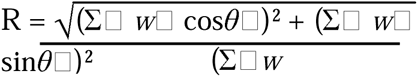 resultant length by:

This routine can be accessed on GitHub, along with the segmented images analyzed for Fig. 6, at the following address:

https://github.com/joshwhiteley/signal-aggregation-comparison

### Lipid Extraction and Data Analysis

Lipids were extracted from WT and CD59 KO 293T cells by the Folch extraction method (44) as follows. CD59 KO and WT 293T cells were grown in DMEM + 10% FBS until ∼80% confluency in one 10cm plate per sample replicate, and medium was changed 2 hours prior to harvesting. Cells were rinsed with PBS, lifted with 0.05% trypsin, spun down for 5min at 200xg, washed with PBS, and resuspended with PBS in a total volume of 200μL. Cells were transferred to glass vials and 2:1 chloroform: methanol was added to a final volume of 4mL. Samples were then agitated for 30 min on an orbital shaker at RT before 800μL water was added, vortexed 1 min, and allowed to stand for 10 minutes before centrifuging for 5min at 1000xg to separate into three phases. The lower phase containing nonpolar lipids was carefully transferred with a glass pipette from the vial into the microtube, then evaporated under nitrogen and stored at -80 °C. Samples were then analyzed via high-resolution LC-MS/MS, and the lipids were identified and quantified with LipidSearch software by the BIDMC-Harvard Mass Spectrometry Facility according to their published protocol (special thanks to Dr. John Asara, Director) (44). Lipid peak data were normalized by percent abundance within each sample. PCA analysis was performed using MetaboAnalyst 5.0. Lipid pathway analysis performed using LIPIDMAPS BioPan software.

## Supporting information

Description of Supplemental Files

## Acknowledgements

We would like to thank Dr. Rodney Tweten (Oklahoma University Health Science Center) for the kind gift of plasmids encoding intermedilysin derivatives and for discussions during the early stages of this work. We would like to thank Stephen Kwok and Alan Parmelee from the Tufts Flow Cytometry Core Facility for assistance in flow cytometry, Dr. Joan Mecsas and her lab for Yersinia strains and discussions, as well as Drs. Marta Gaglia and Katya Heldwein for experimental suggestions. We thank Drs. Efrat Hamami, Kevin Manera, Atish RoyChowdhury and Bixi He for critical review of the manuscript.

## Author Contributions

Kristen Davis: conceptualized and designed the analysis, collected, analyzed, and interpreted the data, writing of original draft, editing of the paper; Kerri Sheahan: conceptualization, constructed CD59 KO cell line, collected preliminary data; Connor Murphy: conceptualization, collected preliminary data; Joshua Whiteley wrote code and performed image analysis; Bree Aldridge oversaw methodology and interpretation of data; Ralph Isberg: conceptualization and design of analysis, funding acquisition, project administration, methodology, interpretation of data, editing and review of manuscript.

## Supplemental Figure Legends

**Figure S1. CD59-KO show a kinetic defect in pore formation.** Noted cell lines adherent to 96-well dishes were challenged with either the YPIIIΔ5 strain (pore-forming competent) or YPIIIΔB (TTSS-defective) in the presence of propidium iodide (PI) in triplicate wells, and mean PI fluorescence was assayed over time using a multiwell fluorometer at 37°C. Shown are mean ± SE for the 3 determinations.

**Figure S2. Pore formation by *Yptb* is T3SS-dependent.** CD59KO HEK 293T cells were transfected with a plasmid expressing human CD59, and stable monoclonal cell lines were obtained with CD59 levels that were similar to endogenously expressed protein (WT1 and WT2) or which were overexpressed (OE1 and OE2). **(A)** Indicated cell lines were challenged with YPIIIΔ*yopHMOJE* (YPIIIΔ5) or YPIIIΔ*yopB* (YPIIIΔB) at MOI = 25 for 2 hrs. Cells were labeled with FITC LIVE/DEAD stain and analyzed by flow cytometry to determine extent of pore formation. Data are geometric mean ± SEM of the fluorescence intensity normalized to WT. Data shown is representative of three individual experiments. **(B)** Pore formation peak data showing WT1, WT2, OE1, and OE2 compared to WT and KO after infection by YPIIIΔ5.

**Figure S3. Treatment by Phospholipase C removes CD59 from cell surface.** WT HEK 293T cells were seeded overnight in 12-well plates. Cells were treated with phospholipase C (PLC) and incubated at 37°C, 5% CO_2_ for 30 mins before harvesting and labeling with FITC-conjugated α-human CD59. Cells were analyzed by flow cytometry for detection of cell surface-localized CD59. Bars note the fluorescence distribution of PLC treated cells.

**Figure S4. siRNA knockdown of CD59 is efficient prior to focal adhesion assay.** WT HeLa cells were transfected with either siCD59 (blue) or siNT (red) to knock down CD59 expression. Assays were performed after 72 total hours of transfection. Cells were probed with FITC-conjugated α-human CD59 and analyzed by flow cytometry for cell surface localization of CD59. Bars note the fluorescence distribution of siCD59 sample.

**Figure S5. Determination of polarity index for perinuclear localization of CtxB-bound GM-1 gangliosides.** Polarization indices for theoretical localization patterns as depicted by three cartoons. **A)** Cartoon showing nucleus fully encompassed by CtxB staining: Polarization Index = 0. **B)** Cartoon showing nucleus abutted by CtxB staining with single-sided bias: Polarization Index = 0.97. **C)** Cartoon showing nucleus abutted by CtxB staining showing randomly dispersed puncta: Polarization Index = 0.14. **D)** Examples of regions of interest (ROI), setting DAPI-stained nucleus as the inside limit of ROI, with the outside limit set as 1μm dilation of the displayed nuclei circumferences. **E)** Six individual cells from the dataset generated in Fig. 5, showing examples of circumferential and polarized CtxB staining. Displayed are the calculated polarization indices (R) for the two states.

**Figure S6.** CD59 knockdown does not change surface levels of GM1 gangliosides. **(A, B)** siNT control cells and siCD59 knockdown cells were probed with FITC-conjugated α-human CD59 **(A)** or immunolabeled for GM1 **(B)** and analyzed by flow cytometry after a 72 hr knockdown.

