## Supplementary material for "Host CD59 Potentiates the Type III Secretion System in *Yersinia pseudotuberculosis*": Description of Supplemental Files

**Description of Additional Supplementary Files**

File Name: pTyrAnalysis1+Exp+ROI.ijm

Description: ImageJ macro file used to identify focal adhesions in images with 3 channels (RGB) using a modified method adapted from the publication "Horzum et. al, MethodsX, 2014"

**File Name: Supplemental_dataset_1**: Data for Fig. 5H to determine the Polarization Index (R) for each of 6 samples displayed in Fig. 5A-F. R determined as described in text.

**File Name: Supplemental_dataset_2.xlsx**

Description: LC-MS/MS peak area values for identified lipid ions of HEK 293T WT cells vs CD59 KO cells. Additional tabs show calculations for:

“PC”: PC species normalized to %sum of total PC species, then normalized peak areas summed by total number of carbons in the acyl chains (Figure 6B).

“PE”: PE species normalized to %sum of total PE species, then normalized peak areas summed by total number of carbons in the acyl chains (Figure 6C).

“SM”: SM species normalized to %sum of total SM species, normalized peak areas summed by total number of carbons in the acyl chains, and normalized peak areas summed by both length and unsaturation level (# of double bonds) for the three most abundant total carbon counts within SM species (34C, 36C, and 42C) (Figure 6D).

“%sumtotallipids”: All individual lipid ion peak areas were normalized by calculating the percentage sum of total lipids. These normalized values were submitted to LIPID MAPS BioPAN for pathway analysis.

“BioPan_Subclass_Rxn_Compare”: Comparison of subclass reaction chains and z-scores of two independent experiments analyzed by LIPID MAPS BioPAN software. Scores with shared directionality for both experiments are represented in Figure 6E.
