## Supplementary figures and images for "Host CD59 Potentiates the Type III Secretion System in *Yersinia pseudotuberculosis*"

### Supplemental Figure S1

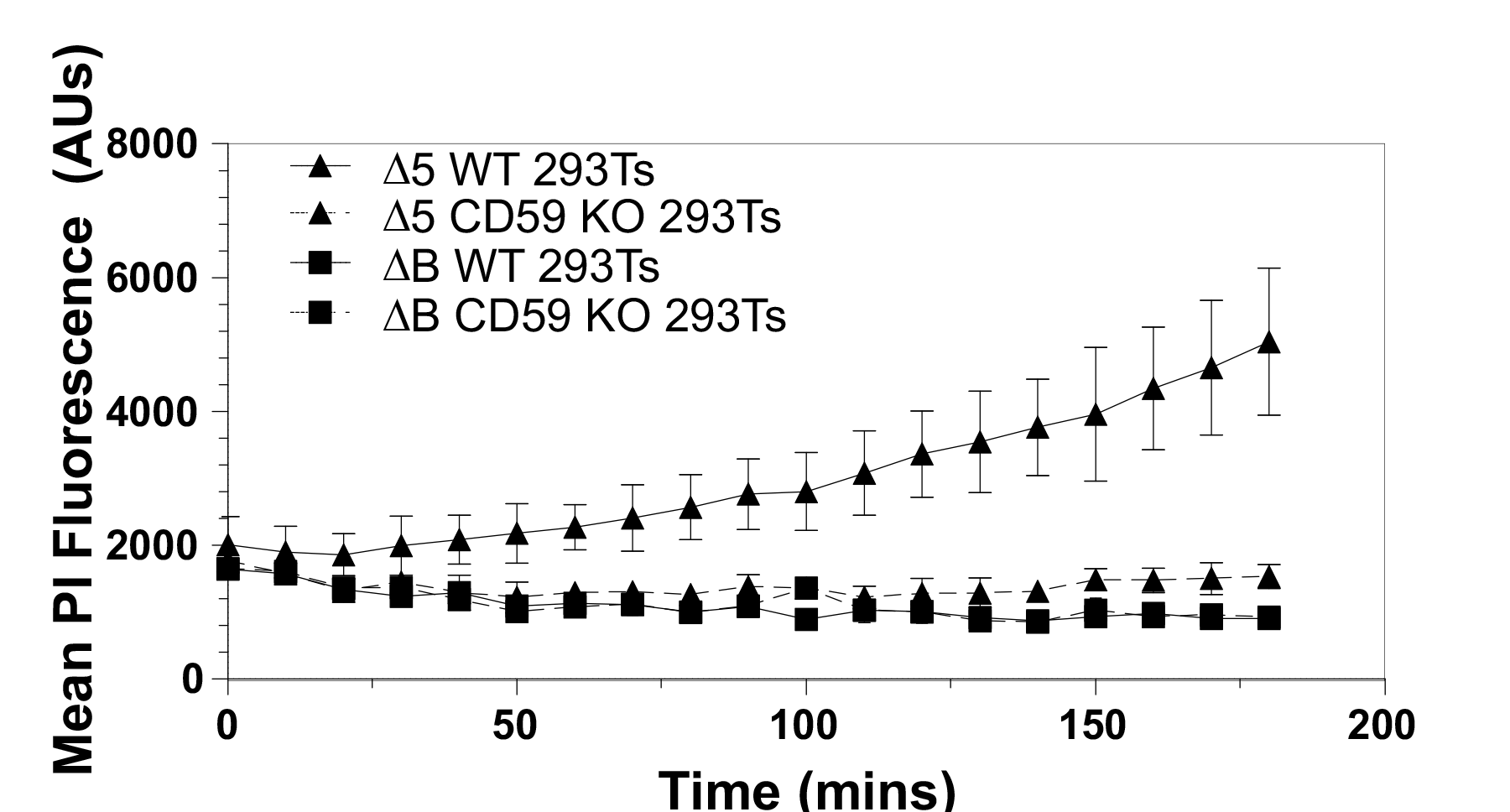

### Supplemental Figure S2

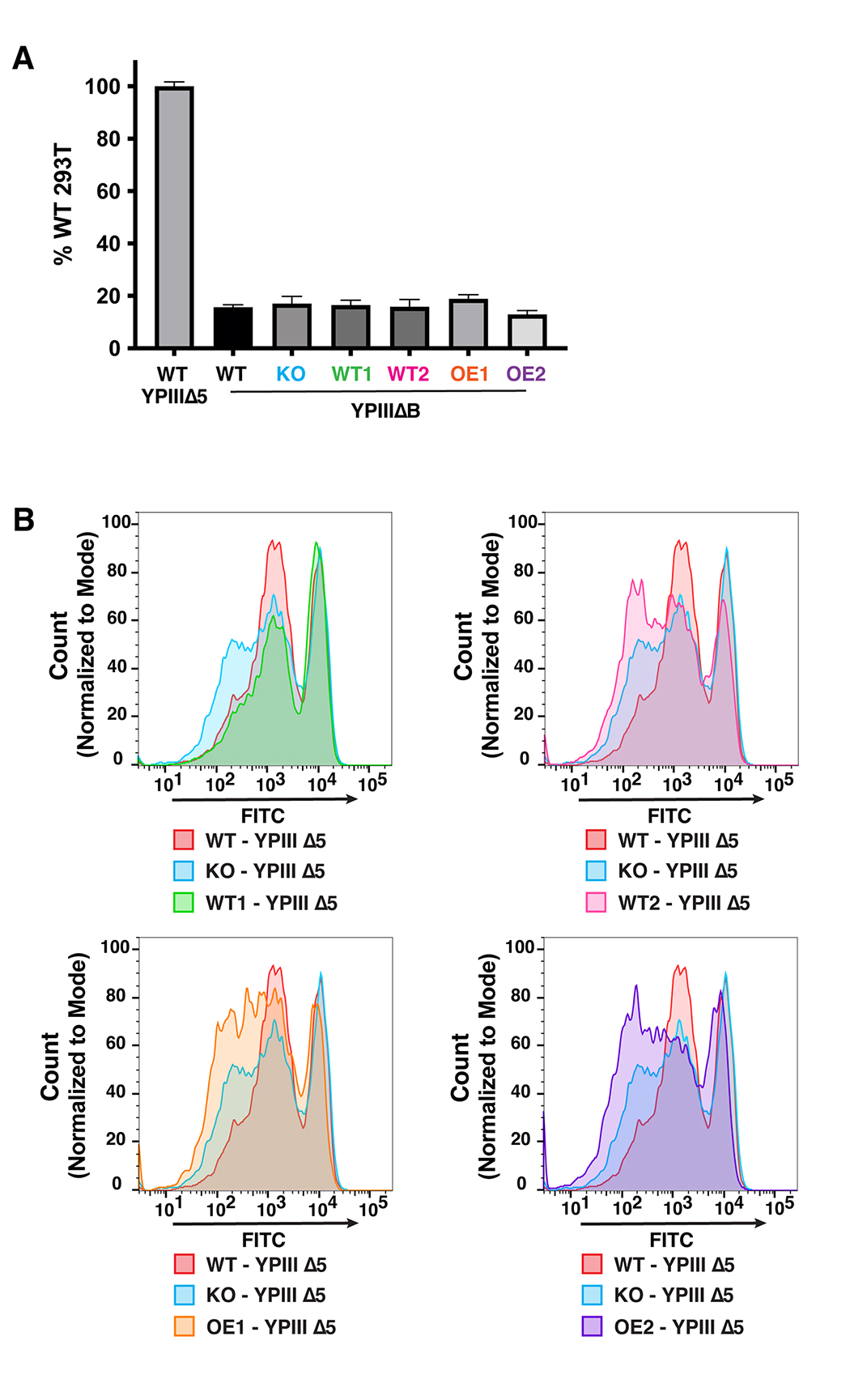

### Supplemental Figure S3

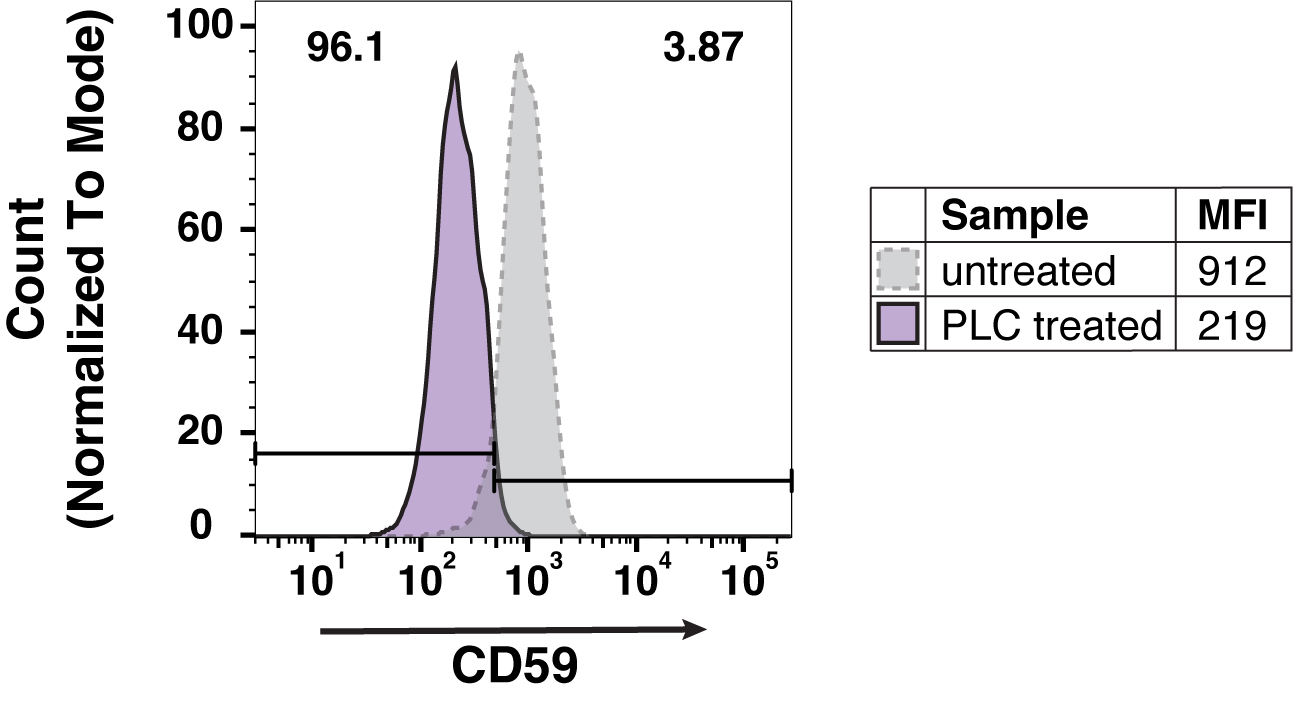

### Supplemental Figure S4

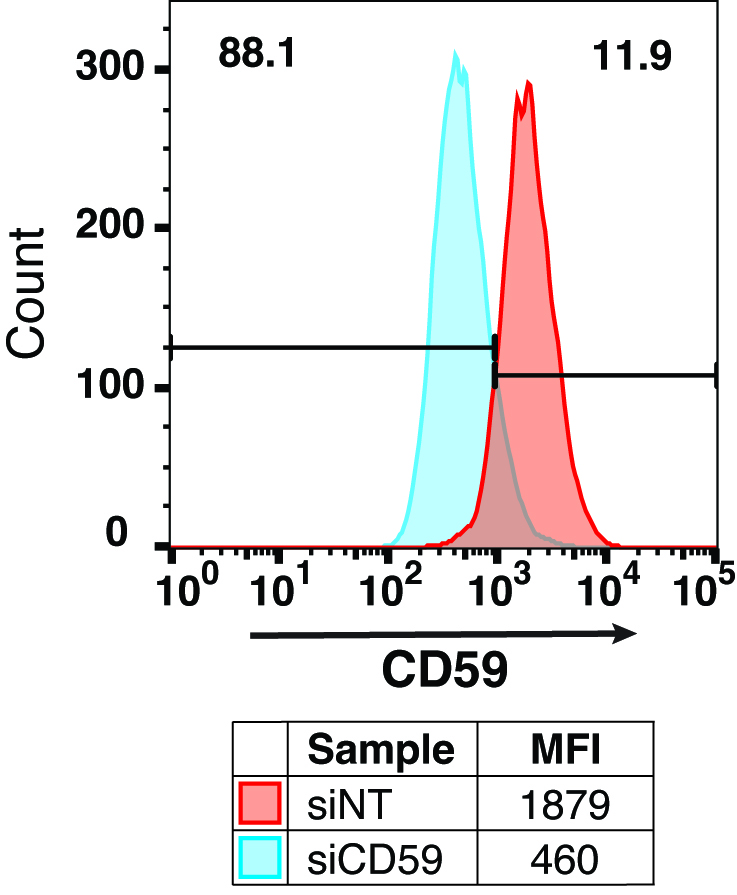

### Supplemental Figure S5

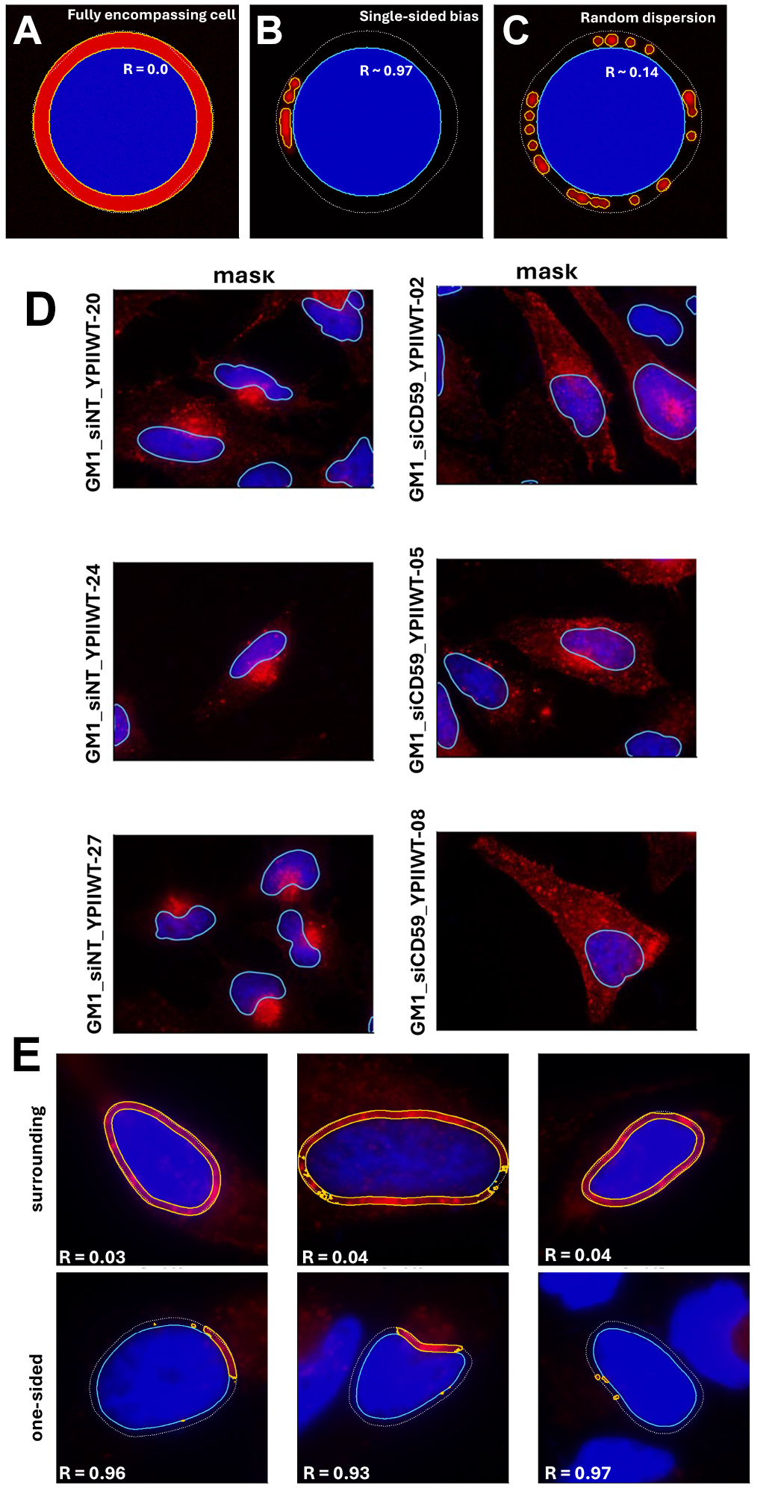

### Supplemental Figure S6

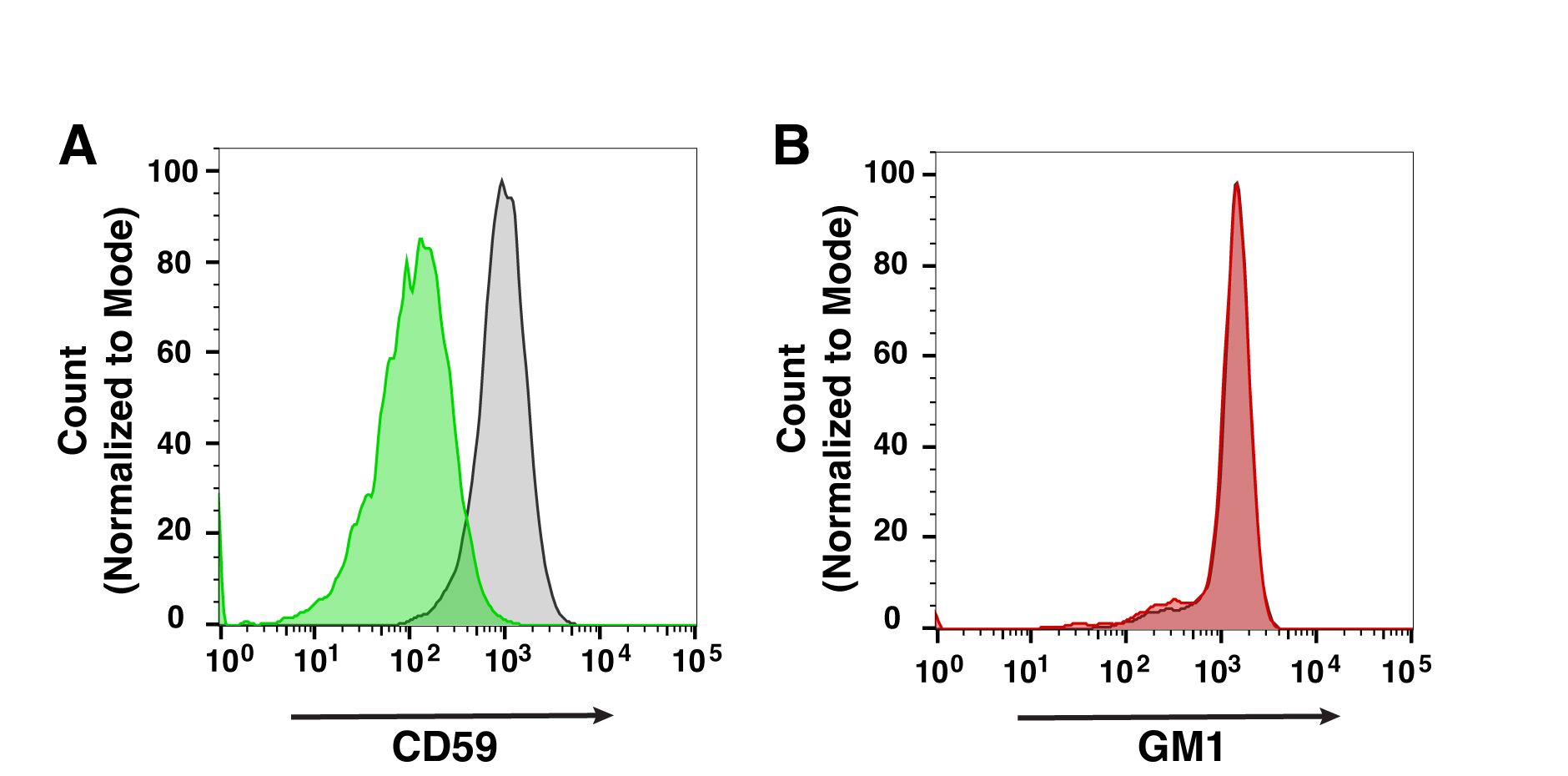
